# Identifying the Levant as a potential contact and interbreeding zone for Neanderthals and modern humans

**DOI:** 10.1101/2025.08.02.668257

**Authors:** Anooshe Kafash, Trine Kellberg Nielsen, Marc Grünig, Jens-Christian Svenning, Masoud Yousefi

## Abstract

Timing of interbreeding between modern humans and Neanderthals has been subject of numerous studies but its geography remains largely unknown. Genetic evidence suggests three different interbreeding events: first in the Marine Isotope Stage (MIS) 7 (∼250 to 200 ka), then in the MIS5 (∼100 to 120 ka) and the final event in the MIS3 (∼60 to 50 ka). Here, we used all known archaeological sites between 60-50 ka associated with Neanderthals and modern human presence and a set of paleoenvironmental data to reconstruct Neanderthals and modern humans’ habitat suitability using the Species Distribution Modeling (SDM) techniques. Assessing geographical overlap between the two species, we identify potential interbreeding zone. We found that the Levant was main potential interbreeding area of the third event. Previous research has identified the Zagros Mountains in Iran as a potential interbreeding zone during the second interbreeding event MIS5 (∼100 to 120 ka). Compiling the results of this study to previous research can help us to better understand the dynamics of modern humans and Neanderthals interbreeding over both time and space. The two potential interbreeding areas have high priority for future research.

## Introduction

*Homo sapiens* (hereafter: modern humans) originated around 300,000 years ago in Africa with warm and savanna-like environments ^1-3^ and then dispersed into Eurasia ^4,5^. One critical topic in human evolutionary research is the dispersal of modern humans out of Africa ^4,6,7^ particularly between 60 and 40 ka ^8,9^. Modern humans encountered *Homo neanderthalensis* (hereafter: Neanderthals) while dispersing across Eurasia which led to interbreeding with them and finally replacing them ^8,10-12^. Unlike modern humans which originated in Africa, Neanderthals, the closest species to modern humans evolving in parallel, first appeared in Europe before 400,000 years ago ^13^. This archaic human species then dispersed in to the Middle East and then Central Asia and Siberia ^14-16^. Like the modern humans, Neanderthals were an opportunistic hominin species, with a diverse diet including nutrition gained from hunting, gathering, fishing and seafood consumption ^17-21^. This archaic human species faced extinction approximately 40,000 years ago ^15,22^.

Studying the extent, timing and geography of interbreeding between modern humans and Neanderthals is fundamental to understand their interactions and impacts on the genomes and biology of the two species ^11,23^. Previous studies significantly increased our knowledge of the two species admixture over the past decade and a half, by identifying three waves of gene flow from Neanderthals to modern humans and vice versa ^11,13,23,24^. The initial wave of interbreeding occurred ∼250 to 200 ka (MIS7), the second wave of interbreeding occurred ∼100 to 120 ka (MIS5) and the third and last interbreeding occurred ∼60 to 50 ka (MIS3). While the timing of admixture between the two human species is known, the geographical area, where the interbreeding may have occurred remains less investigated. To date, only the Zagros Mountains of Iran could be identified as potential interbreeding zone during the MIS5 wave using Ecological Niche Models (ENM)^25^.

We are particularly interested in knowing where interbreeding occurred to identify high priority areas for future studies and as timing of interbreeding is known it is possible to locate potential interbreeding areas ^11,23-25^. Ecological Niche Models (ENM) can be very useful in this matter by reconstructing the distribution of the two species at the estimated time of interbreeding, 60-50 ka. Here, we used all known Neanderthals and modern humans archaeological sites (60-50ka) and a set of high-resolution paleoenvironmental data and employed ENM techniques to reconstruct their global habitat suitability and niche overlap and identify potential interbreeding zones at 60-50 ka ^23,24^. We considered geographic overlaps as a proxy for interbreeding between the two species ^25,26^. We focused on the third interbreeding event because its geography is less known compared to the second interbreeding event ^25^. Since it is claimed that the second wave of interbreeding occurred on Zagros Mountains of the Iranian plateau we predict the third wave also occurred somewhere on the Iranian plateau ^25^.

## Methods

In this study to reconstruct Neanderthals and modern humans ecological niche and their geographic overlap, we used two sets of data. Archaeological site coordinates and environmental predictors characterizing paleo environmental conditions of 60-50ka ^27-30^.

### Neanderthals and modern humans presence data

We used archaeological sites associated with presence of Neanderthals and modern humans across their distribution range ^28,31^. We limited Neanderthals and modern humans presence data to 60 to 50 ka which is the time suggested for the third interbreeding wave ^23^. We included 155 sites for Neanderthals and 39 sites for modern humans in niche modeling. Considering that archaeological sites age estimations are always associated with uncertainties and sometime high temporal intervals we used the mean age of each archaeological site to the time period of the third interbreeding wave, attribute it to 60-50ka ^23^. With this approach our models can predict the two species geographical distribution and overlap with less uncertainties.

### Paleoenvironmental predictors

We extracted five paleoenvironmental predictors (mean temperature (bio01), min temperature (bio05), max temperature (bio06), mean temperature during warm quarter (bio10), mean temperature during cold quarter (bio11), mean precipitation (bio12), mean precipitation during cold quarter (bio19), net primary productivity (npp), and biome) from the Pastclim 1.2 package ^27^. These predictors are available on a millennium time scale at 0.5-degree spatial resolution. We extracted predictors for each year between 60 and 50 ka. To estimate average values for the paleoenvironmental predictors, we used the raster package implemented in R environment. Variable selection was based on the target species ecology ^32^.

We also obtained a set of very high resolution paleoclimatic predictors at 2.5 arc-minutes (4.65 km at the equator) spatial resolution from Oscillayers, which is a dataset of climatic oscillations over Plio-Pleistocene time-scales ^30^. We first downloaded all 19 bioclimatic predictors then calculated a variance inflation factor (VIF; ^33^) using the “vifstep” function in the “usdm” package and only kept those predictors with low collinearity (VIF < 10). Because Oscillayers dataset temporal resolution is 10 ka years we extracted predictors for 60 and 50 ka and then estimated average values for the two time slices using the raster package implemented in R environment

### Ecological niche modeling

We developed two Species Distribution Models (SDMs) with the same approach but different paleoenvironmental predictors, Pastclim and Oscillayers. We used an ensemble of four different niche modeling approaches; Generalized Linear Models (GLM), Generalized Additive Models (GAM), Random Forests (RF), and Maximum Entropy models (Maxent) in the sdm package ^34^. Considering that these methods need pseudo-absences we generated 5000 pseudo- absences by randomly sampling coordinates from the biomes where the two species occurred. We used 75% of the data as the training set and the remaining 25% was used for evaluation. Models were evaluated using Receiver Operating Characteristic (ROC) metric ^35^. AUC ranges from 0 to 1, an AUC value of 0.5 indicates that the performance of the model is not better than random, while values closer to 1.0 indicate better model performance ^35^.

Then we overlapped the two paleodistribution models to identify potential areas for their contact zones in QGIS 3.38.1 (www.qgis.org). To this end, we first converted continues habitat suitability maps to binary (suitable/unsuitable) maps using the averaged values of the maximum test sensitivity plus specificity and maximum training test sensitivity plus specificity thresholds in raster package.

## Results and Discussion

The models developed for Neanderthals (AUC□=□0.936) and modern humans (AUC□=□0.918) performed well according to the AUC model performance metric. Niche models were created based on Neanderthals and modern human archaeological sites present data for 60-50ka (Figure 1). The models developed for Neanderthals and modern humans based on the Pastclim data (Figure 2), show that South and North of Africa, South of Europe and some large patches in Asia were highly suitable for modern humans at 60-50ka. South and West of Europe, coasts of Mediterranean Sea and Black Sea were highly suitable for the presence of Neanderthals. Based on the Oscillayer data South and North of Africa, South of Europe and some large patches in Asia were found to be highly suitable for modern humans at 60-50ka. Europe, coasts of Mediterranean Sea and Black Sea were also highly suitable for the presence of Neanderthals.

**Figure 1.**
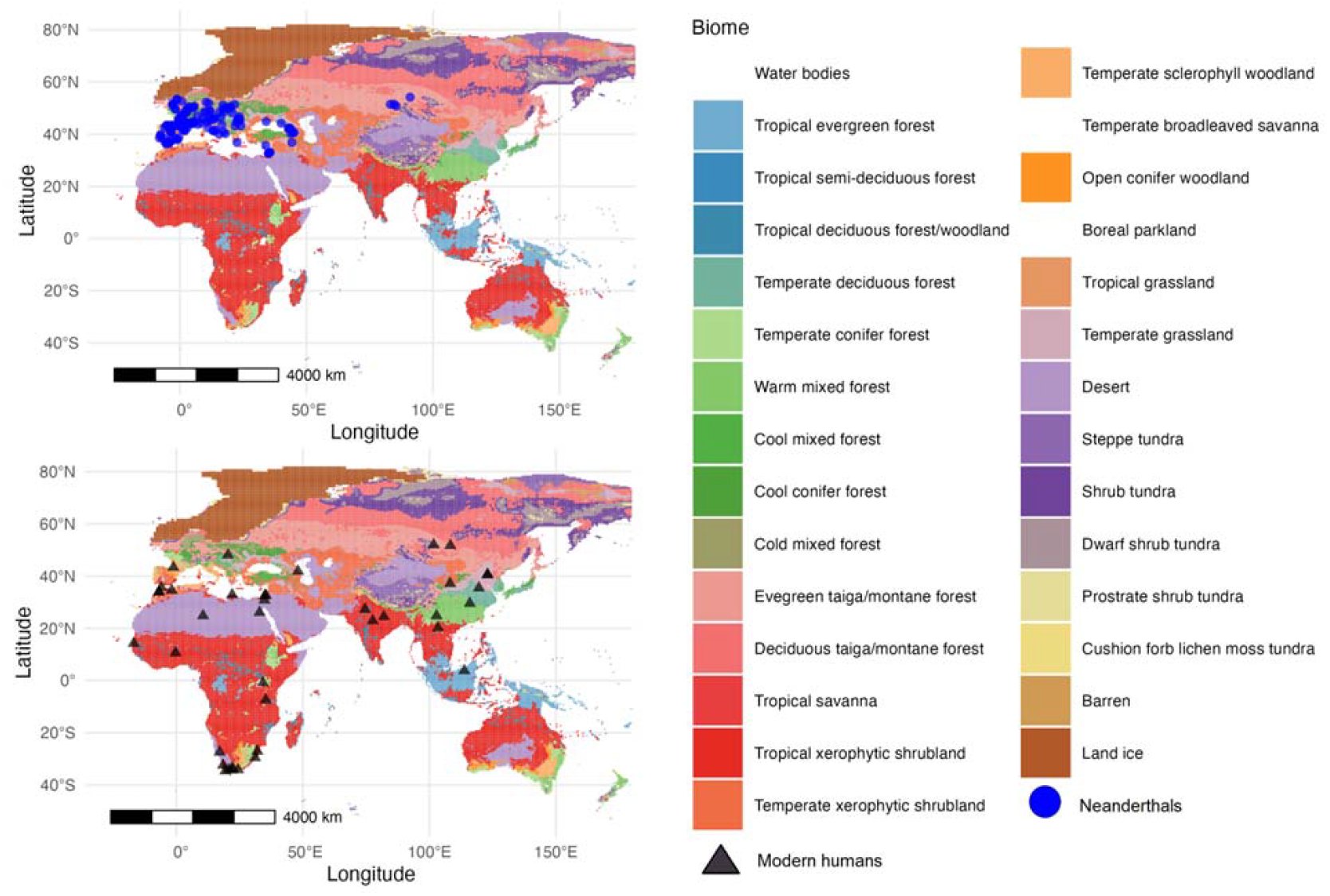
Distribution of Neanderthals and modern humans. Archaeological sites associated with presence of Neanderthals (top) and modern humans (below) in terrestrial biomes. During the 60-50 ka Neanderthals were living within the five different biomes (Figure 1). Thirty one percent of archaeological sites associated with Neanderthals presence are located in Temperate Sclerophyll Woodland biome and 10.8 % in each Temperate Conifer Forest biome and Cool Conifer Forest biome. Modern humans were primarily living in Tropical Xerophytic Shrubland biome (60.7%) and Desert biome (17.9%) was also a major area for their presence during the time. Figures were produced using the R software (https://cran.r-project.org/).

**Figure 2.**
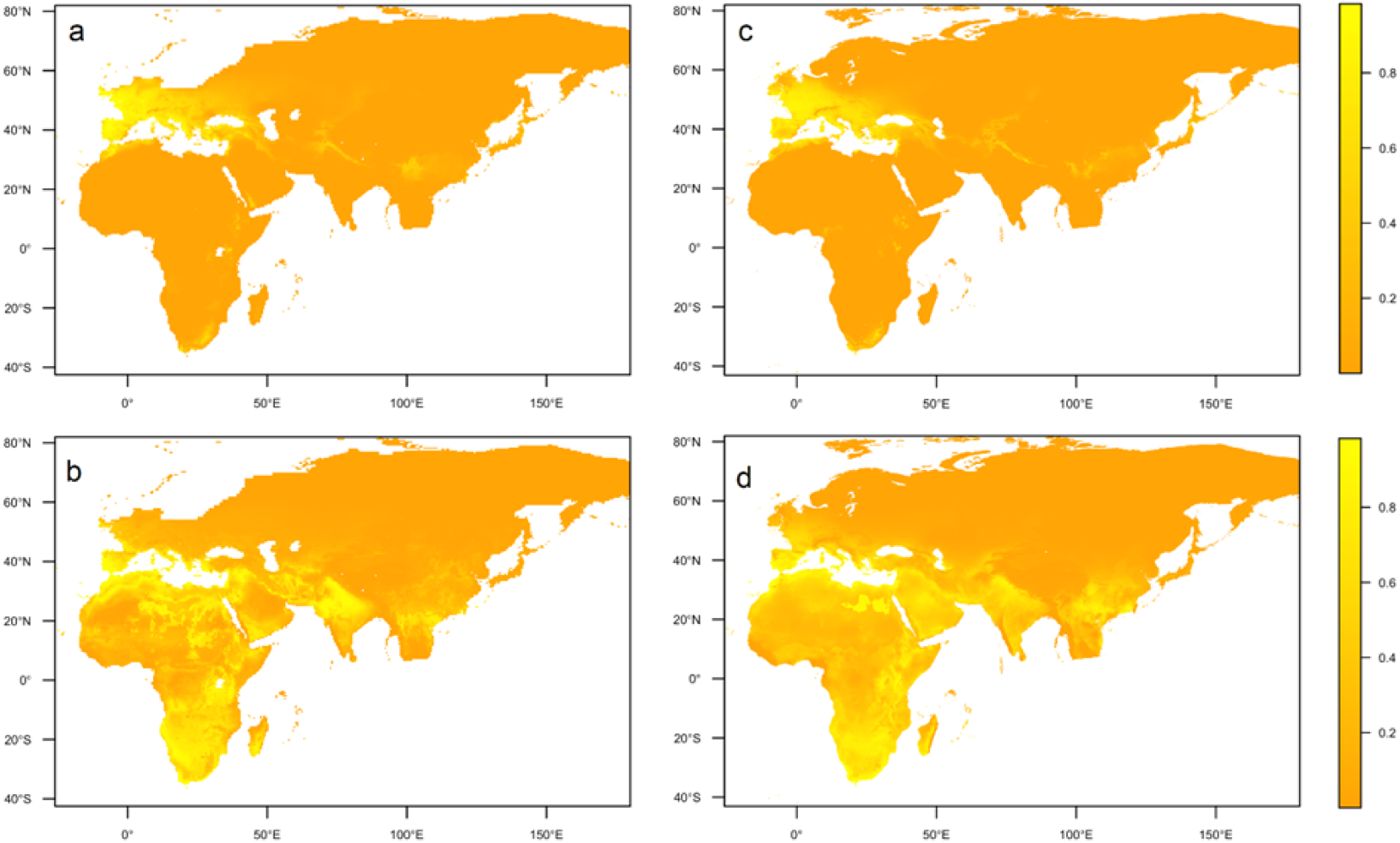
Ecological suitable areas for Neanderthals and modern humans at 50-60ka according to Environmental Niche Modelling. Estimates for Neanderthals (top) and modern humans (below) are based on the Pastclim data (a and b) and Oscillayer data (c and d) at 50 km and ∼5 km spatial resolution respectively. Light yellow areas show suitable range of the two species and orange areas unsuitable range of them. Figures were produced using the R software (https://cran.r-project.org/).

We identified two major niche overlapped areas in which interbreeding took place at 60-50 ka, the Iberian Peninsula and the Levant (Figure 3). The most suitable habitats of Neanderthals were located in Western Europe particularly the Iberian Peninsula at 60 50ka ^32^ where Neanderthals and modern humans live alongside each other in Europe before Neanderthals extinction ^36,37^. This make Western Europe particularly Iberian Peninsula a highly probable area for the two species interbreeding. We also identified some other patches (for example South Africa) where the two human species niche overlap with each other but we argue that because there are not located in the way of out- of-Africa dispersal paths, it makes them weak candidate for such interbreeding event.

**Figure 3.**
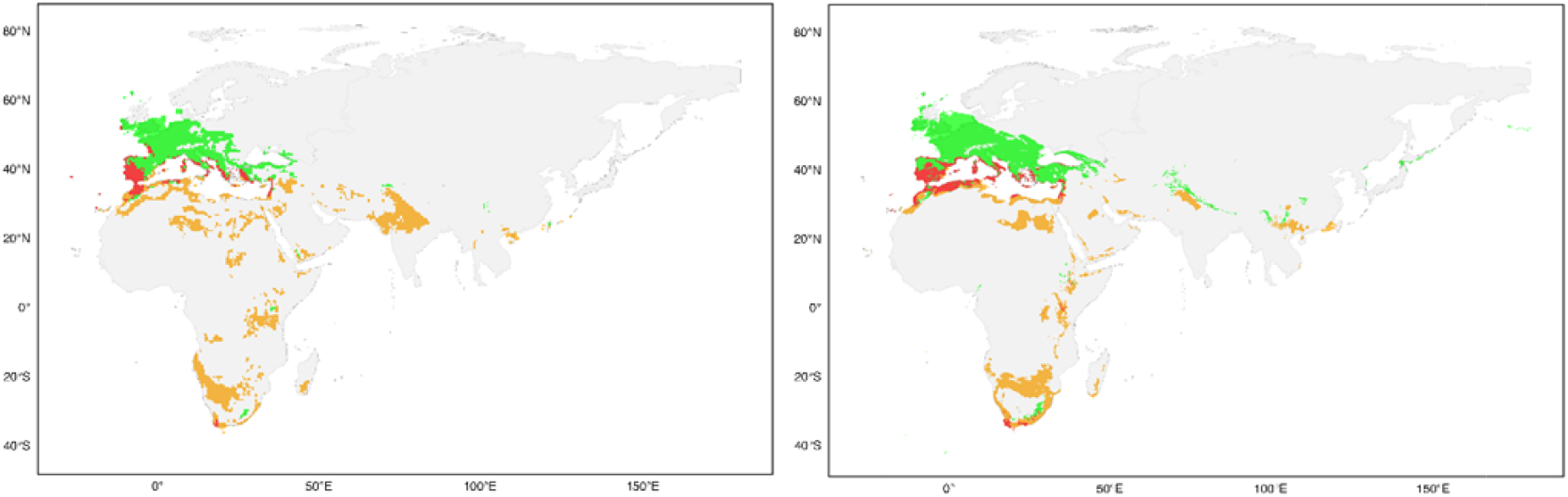
Niches overlap and potential interbreeding areas between Neanderthals and modern humans based on the Pastclim data (right) Oscillayer data (left). Green areas show suitable range of Neanderthals, orange areas suitable range of modern humans and red contact zones and potential interbreeding areas. Figures were produced using the QGIS 3.38.1 (www.qgis.org) and R software (https://cran.r-project.org/).

Potential interbreeding area should be large enough to have high habitat diversity, sustain large population of the two species and enough food resources to support presence of the two species in the same area at the same time^25^. The broad overlap in the Mediterranean region including Asia Minor suggest that other factors have maintained the parapatric distribution of the two species in through the later part of the Middle Pleistocene and early part of the Late Pleistocene.

The Iberian Peninsula which lies in southwestern Europe and is topographically complex and hosts several mountain ranges exceeding 2000–2500 m. The peninsula was identified to be highly suitable for both species and interestingly there is recent evidence suggesting the two species coexisted in the area for 1,400 to 2,900 years ^36,38^.

The Levant overlap area which encompasses the Mediterranean shores of the Middle East, is geographically smaller than the Iberian Peninsula but it is known to be a very important corridor for out-of-Africa scenario ^8,9,39-41^. The Levant linked Africa to Eurasia and is confirmed that Neanderthals and modern humans coexisted in the areas for some time but regarding the small size of the overlap area it can be assumed that small groups of both were living in the area at the same time ^8,39,42^. The two human species may have entered to the region not at the same time as Neanderthals were a resident population in the Levant before modern humans (Upper Palaeolithic) populated the region ^39^.

The two geographic regions are probably areas in which modern humans first encountered Neanderthals while migrating out-of-Africa. The Levant is one of the most known and highlighted corridors for before modern humans (Upper Palaeolithic) migrating out-of-Africa ^9,39-41^. While the overall geographic area of the Levant is small compared to other corridors like Arabian Peninsula but it played a very critical role in modern humans’ migration out-of-Africa. More importantly Neanderthals and modern humans coexisted in this corridor as it is shown by our models and archaeological evidence.

Our habitat suitability models of 60-50 ka for modern humans show a connection between southernmost Spain and northwesternmost Africa (Strait of Gibraltar) which could be a possible route for rapid expansion of modern humans across Europe 60-50ka, the Southwest European out-of-Africa corridor. During the time sea level was lower and a connection between the two continents is possible. This potential dispersal route is also matched with the ideas of rapid dispersal of modern humans along coastal landscape ^43^. Areas close to the Strait of Gibraltar in both continents have high priority for future investigations. More research in these areas could eventually paint a more detailed picture of the two species coexistence, interactions and interbreeding. It should be noted that there (the Strait of Gibraltar) always was a sea barrier which prohibited exchange of other non-flying terrestrial biota between Africa and Europe. But we think that the more conventional alternative corridor is that modern humans’ immigration into the region came from the east, via the Balkans and Italy.

Paleo-environmental data sets are increasing in recent years providing important opportunities to study past distribution of extinct and extant species including human species ^27,30,44,45^. Here we applied ecological niche modeling methods to locate potential areas for the last wave of Neanderthals and modern humans interbreeding. While our approach can be used to uncover potential areas for the first interbreeding event ^23^, but lack of enough archaeological sites associated with presence of the two species for the first interbreeding event makes us unable to build robust niche models. We recommend the two overlap areas, the Iberian Peninsula and the Levant coincide with the archaeological and genetic data, to be investigated and excavated by field archaeologists as these potential interbreeding areas hold significant information which can increase our understanding of the two species interaction and interbreeding.

Here we identified two overlap area for the Neanderthals and modern humans at 60-50 ka, the Iberian Peninsula and the Levant. Identification of these interbreeding areas using ecological niche models can be supported with archaeological and fossil data ^36,39^. We recommend the potential interbreeding areas for future archaeological and paleoanthropological studies to collect more support for the two areas. The newly identified Southwest European out-of-Africa corridor was not recognized in previous studies but has potentials to receive attention in future research.

## Data availability

All data necessary to evaluate the conclusions of this study are provided within the main text or in the references cited herein.

## Acknowledgments

This study is part of the NeanderEDGE project, which is funded by the Independent Research Fund Denmark (Case Number 9062-00027B). The Center for Ecological Dynamics in a Novel Biosphere (ECONOVO), funded by Danish National Research Foundation (grant DNRF173).

## Author contributions

A.K. conceived and designed this study. A.K. collected distribution data, A.K. and M.Y. performed statistical analyses. A.K. wrote the first draft of manuscript. T.K.N. supervised the study. A.K., T.K.N., M.G., J.C.S. and M.Y. read, revised, and approved the final manuscript.

## References

1 Stringer, C. & Galway-Witham, J. On the origin of our species. Nature 546, 212–214, doi:10.1038/546212a (2017).

2 Hublin, J.-J. et al. New fossils from Jebel Irhoud, Morocco and the pan-African origin of Homo sapiens. Nature 546, 289–292, doi:10.1038/nature22336 (2017).

3 Stringer, C. Why we are not all multiregionalists now. Trends in Ecology & Evolution 29, 248–251, doi:10.1016/j.tree.2014.03.001 (2014).

4 Groucutt, H. S. et al. Multiple hominin dispersals into Southwest Asia over the past 400,000 years. Nature 597, 376–380, doi:10.1038/s41586-021-03863-y (2021).

5 Beyer, R. M., Krapp, M., Eriksson, A. & Manica, A. Climatic windows for human migration out of Africa in the past 300,000 years. Nature Communications 12, 4889, doi:10.1038/s41467-021-24779-1 (2021).

6 Stringer, C. Coasting out of Africa. Nature 405, 25–27, doi:10.1038/35011166 (2000).

7 Groucutt, H. S. et al. Rethinking the dispersal of Homo sapiens out of Africa. Evolutionary Anthropology: Issues, News, and Reviews 24, 149–164, doi:10.1002/evan.21455 (2015).

8 Hershkovitz, I. et al. Levantine cranium from Manot Cave (Israel) foreshadows the first European modern humans. Nature 520, 216–219, doi:10.1038/nature14134 (2015).

9 Bosch, M. D. et al. New chronology for Ksâr ‘Akil (Lebanon) supports Levantine route of modern human dispersal into Europe. Proceedings of the National Academy of Sciences 112, 7683–7688, doi:10.1073/pnas.1501529112 (2015).

10 Larena, M. et al. Philippine Ayta possess the highest level of Denisovan ancestry in the world. Current Biology 31, 4219-4230.e4210, doi:10.1016/j.cub.2021.07.022 (2021).

11 Villanea, F. A. & Schraiber, J. G. Multiple episodes of interbreeding between Neanderthal and modern humans. Nature Ecology & Evolution 3, 39–44, doi:10.1038/s41559-018-0735-8 (2019).

12 Li, L., Comi, T. J., Bierman, R.F. & Akey, J. M. Recurrent gene flow between Neanderthals and modern humans over the past 200,000 years. Science 385, eadi1768, doi:10.1126/science.adi1768.

13 Green, R. E. et al. A Draft Sequence of the Neanderthals Genome. Science 328, 710–722, doi:10.1126/science.1188021 (2010).

14 Krause, J. et al. Neanderthals in central Asia and Siberia. Nature 449, 902–904, doi:10.1038/nature06193 (2007).

15 Higham, T. et al. The timing and spatiotemporal patterning of Neanderthal disappearance. Nature 512, 306–309, doi:10.1038/nature13621 (2014).

16 Kolobova, K. A. et al. Archaeological evidence for two separate dispersals of Neanderthals into southern Siberia. Proceedings of the National Academy of Sciences 117, 2879–2885, doi:10.1073/pnas.1918047117 (2020).

17 Zilhão, J. et al. Last Interglacial Iberian Neanderthals as fisher-hunter-gatherers. Science 367, eaaz7943, doi:10.1126/science.aaz7943 (2020).

18 Naito, Y. I. et al. Ecological niche of Neanderthals from Spy Cave revealed by nitrogen isotopes of individual amino acids in collagen. Journal of Human Evolution 93, 82–90, doi:10.1016/j.jhevol.2016.01.009 (2016).

19 Stringer, C. B. et al. Neanderthal exploitation of marine mammals in Gibraltar. Proceedings of the National Academy of Sciences 105, 14319–14324, doi:10.1073/pnas.0805474105 (2008).

20 El Zaatari, S., Grine, F. E., Ungar, P. S. & Hublin, J.-J. Neanderthals versus Modern Human Dietary Responses to Climatic Fluctuations. PLOS ONE 11, e0153277, doi:10.1371/journal.pone.0153277 (2016).

21 Roberts, P. & Stewart, B. A. Defining the ‘generalist specialist’ niche for Pleistocene Homo sapiens. Nature Human Behaviour 2, 542–550, doi:10.1038/s41562-018-0394-4 (2018).

22 Devièse, T. et al. Reevaluating the timing of Neanderthal disappearance in Northwest Europe. Proceedings of the National Academy of Sciences 118, e2022466118, doi:10.1073/pnas.2022466118 (2021).

23 Li, L., Comi, T. J., Bierman, R. F. & Akey, J. M. Recurrent gene flow between Neanderthals and modern humans over the past 200,000 years. Science 385, eadi1768, doi:doi:10.1126/science.adi1768 (2024).

24 Sankararaman, S., Patterson, N., Li, H., Pääbo, S. & Reich, D. The Date of Interbreeding between Neanderthals and Modern Humans. PLOS Genetics 8, e1002947, doi:10.1371/journal.pgen.1002947 (2012).

25 Heydari-Guran, S., Yosefi, M., Kafash, A. & Ghasidian, E. Reconstructing Contact and Potential Interbreeding Geographical Zone for Neanderthals and Anatomically Modern Humans. (2024).

26 Ruan, J. et al. Climate shifts orchestrated hominin interbreeding events across Eurasia. Science 381, 699–704, doi:10.1126/science.add4459 (2023).

27 Leonardi, M., Hallett, E. Y., Beyer, R., Krapp, M. & Manica, A. pastclim 1.2: an R package to easily access and use paleoclimatic reconstructions. Ecography 2023, e06481, doi:10.1111/ecog.06481 (2023).

28 Raia, P. et al. Past Extinctions of <em>Homo</em> Species Coincided with Increased Vulnerability to Climatic Change. One Earth 3, 480–490, doi:10.1016/j.oneear.2020.09.007 (2020).

29 Timmermann, A. Quantifying the potential causes of Neanderthal extinction: Abrupt climate change versus competition and interbreeding. Quaternary Science Reviews 238, 106331, doi:10.1016/j.quascirev.2020.106331 (2020).

30 Gamisch, A. Oscillayers: A dataset for the study of climatic oscillations over Plio-Pleistocene time-scales at high spatial-temporal resolution. Global Ecology and Biogeography 28, 1552–1560, doi:10.1111/geb.12979 (2019).

31 Timmermann, A. et al. Climate effects on archaic human habitats and species successions. Nature 604, 495–501, doi:10.1038/s41586-022-04600-9 (2022).

32 Yaworsky, P. M., Nielsen, E. S. & Nielsen, T. K. The Neanderthal niche space of Western Eurasia 145 ka to 30 ka ago. Scientific Reports 14, 7788, doi:10.1038/s41598-024-57490-4 (2024).

33 Quinn, G. P. & Keough, M. J. Experimental Design and Data Analysis for Biologists. (Cambridge University Press, 2002).

34 Naimi, B. & Araújo, M. B. sdm: a reproducible and extensible R platform for species distribution modelling. Ecography 39, 368–375, doi:10.1111/ecog.01881 (2016).

35 Fielding, A. H. & Bell, J. F. A review of methods for the assessment of prediction errors in conservation presence/absence models. Environmental Conservation 24, 38–49, doi:10.1017/S0376892997000088 (1997).

36 Djakovic, I., Key, A. & Soressi, M. Optimal linear estimation models predict 1400– 2900 years of overlap between Homo sapiens and Neanderthals prior to their disappearance from France and northern Spain. Scientific Reports 12, 15000, doi:10.1038/s41598-022-19162-z (2022).

37 Hublin, J.-J. et al. Initial Upper Palaeolithic Homo sapiens from Bacho Kiro Cave, Bulgaria. Nature 581, 299–302, doi:10.1038/s41586-020-2259-z (2020).

38 Sala, N. et al. Nobody’s land? The oldest evidence of early Upper Paleolithic settlements in inland Iberia. Science Advances 10, eado3807, doi:10.1126/sciadv.ado3807.

39 Been, E. et al. The first Neanderthal remains from an open-air Middle Palaeolithic site in the Levant. Scientific Reports 7, 2958, doi:10.1038/s41598-017-03025-z (2017).

40 Luis, J. R. et al. The Levant versus the Horn of Africa: evidence for bidirectional corridors of human migrations. Am J Hum Genet 74, 532–544, doi:10.1086/382286 (2004).

41 Abbas, M. et al. Human dispersals out of Africa via the Levant. Science Advances 9, eadi6838, doi:doi:10.1126/sciadv.adi6838 (2023).

42 Valladas, H. et al. Thermoluminescence dates for the Neanderthal burial site at Kebara in Israel. Nature 330, 159–160, doi:10.1038/330159a0 (1987).

43 Mirazón Lahr, M. & Foley, R. A. Towards a theory of modern human origins: Geography, demography, and diversity in recent human evolution. American Journal of Physical Anthropology 107, 137–176, doi:10.1002/(SICI)1096-8644(1998)107:27+<137::AID-AJPA6>3.0.CO;2-Q (1998).

44 Beyer, R. M., Krapp, M. & Manica, A. High-resolution terrestrial climate, bioclimate and vegetation for the last 120,000 years. Scientific Data 7, 236, doi:10.1038/s41597-020-0552-1 (2020).

45 Brown, J. L., Hill, D. J., Dolan, A. M., Carnaval, A. C. & Haywood, A. M. PaleoClim, high spatial resolution paleoclimate surfaces for global land areas. Scientific Data 5, 180254, doi:10.1038/sdata.2018.254 (2018).

